# Effect of exportin 1/XPO1 nuclear export pathway inhibition on coronavirus replication

**DOI:** 10.1101/2023.02.09.527884

**Authors:** Masmudur M. Rahman, Bereket Estifanos, Honor L. Glenn, Ami D. Gutierrez-Jensen, Karen Kibler, Yize Li, Bertram Jacobs, Grant McFadden, Brenda G. Hogue

## Abstract

Nucleocytoplasmic transport of proteins using XPO1 (exportin 1) plays a vital role in cell proliferation and survival. Many viruses also exploit this pathway to promote infection and replication. Thus, inhibiting the XPO1-mediated nuclear export pathway with selective inhibitors has a diverse effect on virus replication by regulating antiviral, proviral, and anti-inflammatory pathways. The XPO1 inhibitor, Selinexor, is an FDA-approved anticancer drug predicted to have antiviral or proviral functions against viruses. Here, we observed that pretreatment of cultured cell lines from human or mouse origin with nuclear export inhibitor Selinexor significantly enhanced protein expression and replication of Mouse Hepatitis Virus (MHV), a mouse coronavirus. Knockdown of cellular XPO1 protein expression also significantly enhanced the replication of MHV in human cells. However, for SARS-CoV-2, selinexor treatment had diverse effects on virus replication in different cell lines. These results indicate that XPO1-mediated nuclear export pathway inhibition might affect coronavirus replication depending on cell types and virus origin.

## Introduction

The nucleocytoplasmic transport of proteins and other molecules is a highly regulated cellular process. Many nuclear export (called exportins) and import (called importins) proteins are involved in this process by exploiting nuclear pore complexes that form proteinaceous channels in the nuclear envelope (1, 2). Exportin 1 (XPO1), also known as CRM1 (chromosome region maintenance 1), is one of the major nuclear export proteins, transporting hundreds of cellular cargo proteins involved in diverse cellular processes such as transcription, translation, cellular growth, differentiation, and mediating inflammatory responses that include antiviral pathways (3, 4). Thus, direct inhibition of XPO1 function has been explored as a potential antiviral and anticancer therapeutic target (5–8). Various natural or chemically synthesized small molecules that bind to XPO1 and block the export of XPO1 cargo proteins from the nucleus to the cytoplasm have been developed (9, 10). Collectively these molecules are known as selective inhibitors of nuclear export (SINEs). SINEs have been shown to have anticancer activity against diverse types of human cancers and at least one modified drug version with increased safety profile, called Selinexor, has been approved by the FDA for treating multiple myeloma and diffuse large B-cell lymphoma (11, 12). SINEs are also reported in the literature to have antiviral activity against RNA viruses like influenza and respiratory syncytial virus (RSV) that causes respiratory infections due to the blocking of key cellular processes and virus-mediated hijacking of the nucleocytoplasmic transport process (6, 13, 14).

Coronaviruses are enveloped, positive-sense RNA viruses that are known to infect humans and animals. At least seven coronaviruses are known to infect humans. Coronaviruses 229E, NL63, OC43, and HKU1 infect people around the world to cause the common cold (15). Other coronaviruses, MERS-CoV, SARS-CoV, and SARS-CoV-2, are responsible for major disease outbreaks and cause high mortality (16). In nature, bats are known to be primary reservoirs of many coronaviruses, from where viruses spill over to intermediate animals such as palm civet, pangolins, rodents, camels, pigs, cattle where viruses evolve and then spillover to humans, causing mild to severe disease (17, 18). The rapid emergence of different variants of SARS-CoV-2 indicates that the viral genome acquire multiple mutations during replication (19, 20). In this context, it is crucial to study the host factors and compounds that can enhance or reduce replication of SARS-CoV-2 (21–25) .

We previously reported that XPO1 inhibitors, including leptomycin B (LMB) and Selinexor, enhanced the replication of oncolytic myxoma virus (MYXV), a member of the leporipoxvirus genus of *poxviridae,* in diverse types of cultured human cancer cells where intrinsic cellular pathways restrict virus replication (26). Selinexor treatment reduced the formation of cytoplasmic DHX9 antiviral granules, which are involved in lowering MYXV late protein synthesis and replication (26). These findings led us to investigate further the role of Selinexor-mediated inhibition of XPO1 nuclear export pathway on other viruses such as coronavirus.

Here, we show that Selinexor treatment can enhance coronavirus replication in a cell type specific manner. Mouse Hepatitis Virus (MHV) replication was enhanced in both murine and human cell lines. Furthermore, the targeted knockdown of XPO1 using siRNA also enhanced coronavirus replication, suggesting the pro-coronavirus drug action is indeed related to the expected nuclear export pathway target. However, SARS-CoV-2 replication remained unaffected in most of the cell lines that were tested. These observations indicate that unlike other viruses, inhibition of nuclear export pathway can have different effect on coronavirus replication in different cell types in vitro.

## Results

### Nuclear export inhibitors enhance gene expression and replication of coronavirus in murine L2 cells

Murine L2 cells are naturally infected by the mouse hepatitis virus (MHV). We first tested whether inhibition of XPO1-mediated nuclear export pathway in L2 cells by different SINEs can alter the replication of MHV. Since these compounds including Leptomycin B (LMB) are toxic to the cells above 1µM concentration, we used lower concentrations that had minimal or no cytotoxicity to the cells after 24 hours. To our surprise, all the tested SINEs, including LMB, significantly enhanced MHV replication with a concentration of 0.1µM and 0.01µM (Fig 1 A-C). Compared to untreated cells, treatment with XPO1 inhibitors Ratjadone enhanced virus titer almost 1.0 log, whereas Anguinomycin and LMB enhanced virus titer about 0.7 log. Based on these results we next tested the FDA approved anticancer nuclear export inhibitor Selinexor in our subsequent experiments. We monitored and measured the viability of L2 cells in response to treatment with different concentrations of Selinexor (Fig 2A). L2 cell viability was reduced only after treatment with a Selinexor concentration of 5µM or above but had minimal or no effect with a concentration of 1 µM or less. To test the effect of Selinexor on MHV replication, L2 cells were first pretreated with different concentrations of Selinexor for one hour and infected with a GFP-expressing MHV A59 (rA59/S_MHV-2_-EGFP) to monitor the level of GFP expression in the infected cells (27–29). Infection with this MHV exhibits reduced cell fusion since the virus expresses a less fusogenic form of the spike protein and thus allowed counting the number of GFP positive cells. We observed increased % of GFP expressing cells when pretreated with Selinexor. (Fig 2B). At 16 hpi, compared to untreated cells, we observed significantly increased % of GFP positive cells after treatment with more than 1µM selinexor. However, after 24 hpi, even 0.0001 µM selinexor treatment showed significantly increased % of GFP expressing cells (Fig 2C). To assess the number of viruses produced in the presence of Selinexor, L2 cells were treated with different concentration of Selinexor and infected with another GFP expressing MHV A59 (MHVE-GFP) that expresses a highly fusogenic form of spike and readily forms plaques. After 24h post infection, the infected cells and supernatant were collected to titer the progeny virus formation by plaque assay. Virus titration results show a significantly enhanced number of viruses in the cells pretreated with Selinexor between 1 and 10 µM concentration (Fig 2D). These results confirm that Selinexor enhances MHV replication and progeny virus formation in naturally permissive L2 cells.

**Figure 1:**
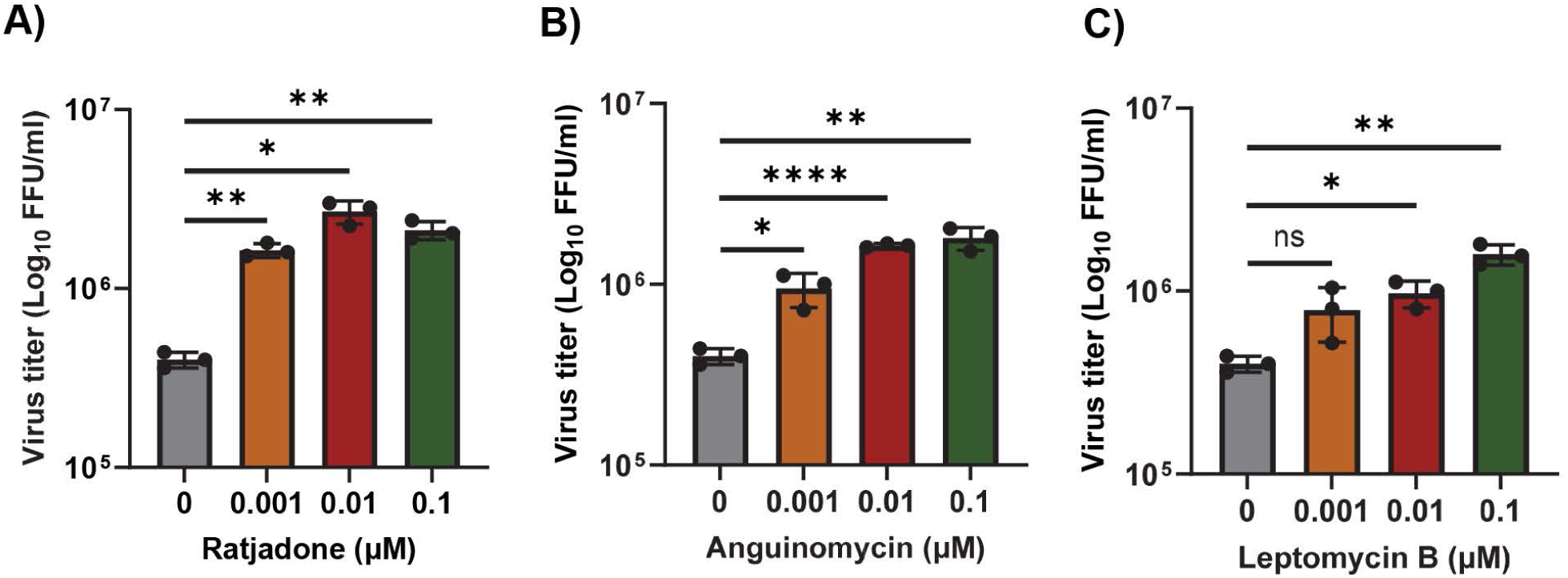
Nuclear export inhibitors enhance the replication of MHV in L2 cells. L2 cells were plated in 24 well plates, treated with the indicated concentration of nuclear export inhibitors: A) Ratjadone, B) Anguinomycin, and C) Leptomycin B for 1h and infected with MHVE-GFP virus with a MOI of 0.01. Virus replication was measured by plaque assay from the total number of viruses in the cells and supernatant 24h post infection. Data represents ± SD and n=3. Statistically significant differences in comparison to 24 hpi infection are indicated. ^ns^ *P* > 0.05, * *P* < 0.05, ** *P* < 0.01, *** *P* < 0.001, **** *P* < 0.0001.

**Figure 2:**
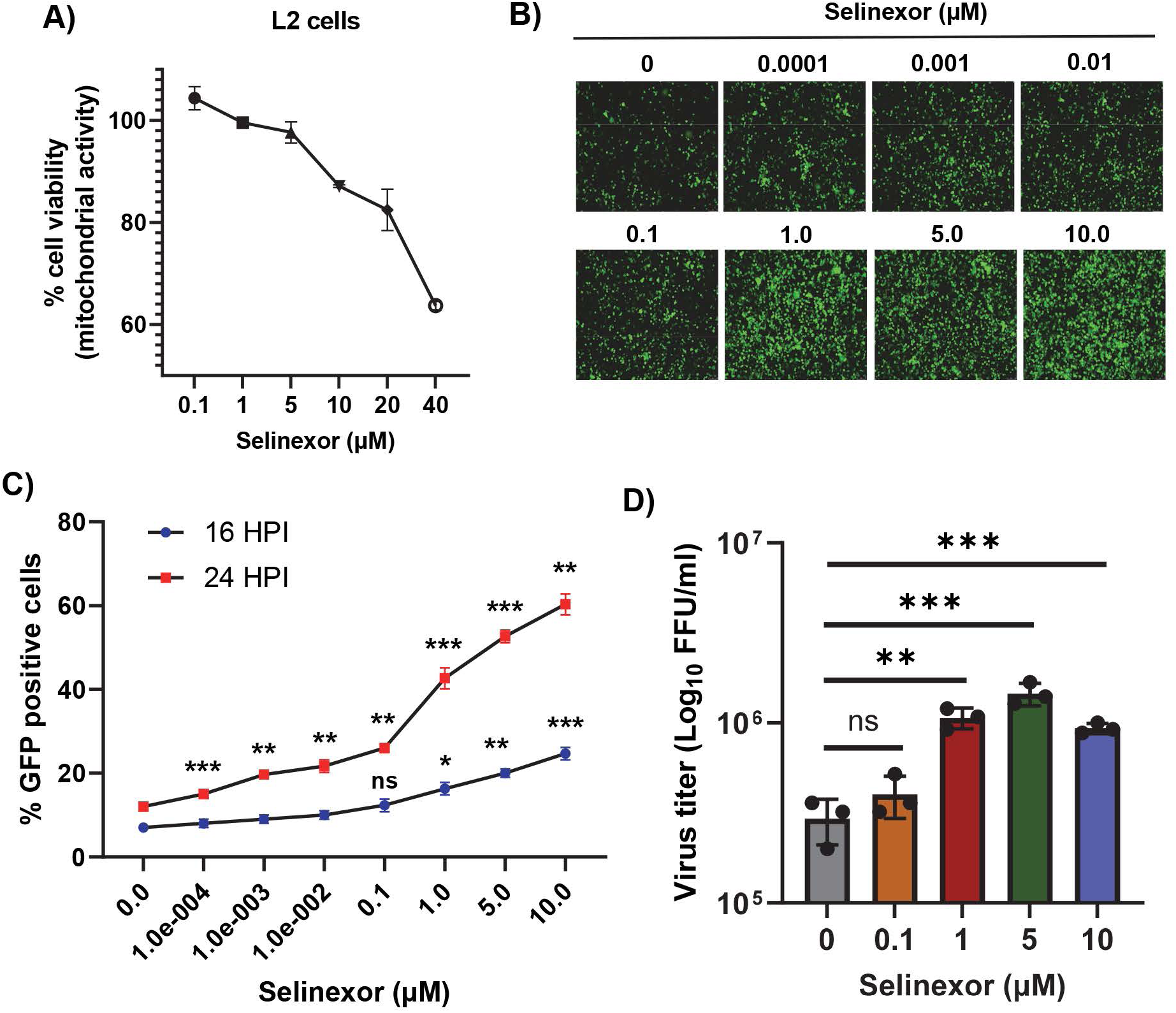
Selinexor enhances the replication of MHV in L2 cells. A) Effect of Selinexor on L2 cells viability. Cells were plated in 96 well plates and treated with different concentration of Selinexor and cell viability was measured after 48h. B-D) Effect of Selinexor on MHV virus gene expression and replication. Cells were plated in 24 well plates, treated with the indicated concentration of Selinexor for 1h and infected with rA59/S_MHV-2_-EGFP or MHVE-GFP virus with a MOI of 0.01. B) Fluorescence images were taken 24h post infection with rA59/S_MHV-2_-EGFP; C) percent of GFP positive cells were counted using Countess II cell counter at different time points after infection with rA59/S_MHV-2_-EGFP; D) MHVE-GFP virus replication was measured by plaque assay from the total number of viruses in the cells and supernatant 24h post infection. Data represents ± SD and n=3. Statistically significant differences in comparison to control are indicated. ^ns^ *P* > 0.05, * *P* < 0.05, ** *P* < 0.01, *** *P* < 0.001.

### Selinexor enhances gene expression and replication of coronavirus in human cells

To further confirm the effects of Selinexor on MHV replication are not cell specific, we used human cell lines expressing MHV receptor (MHVR) carcinoembryonic antigen-related cell adhesion molecule 1 (CECAM1), HeLa-MHVR (human HeLa cell line expressing MHVR) (30) and A549-CECAM/MHVR (human A549 cell line expressing MHVR) (31). Both cell lines were first treated with different concentrations of Selinexor to test cell viability (Fig 3A and 4A). Unlike L2 cells, Selinexor concentrations above 1µM significantly reduced viability of the human cells. However, with a concentration of less than 0.1µM, we observed minimal or no reduction in cell viability. To assess whether Selinexor enhances MHV replication in the human cell lines, both were pretreated with different concentrations of Selinexor for one hour and infected with a GFP-expressing MHV, rA59/S_MHV-2_-EGFP to monitor GFP expression in the infected cells. We observed enhanced GFP expressions when pretreated with Selinexor at a concentration of 0.01 µM or more (Fig 3B). Enhanced GFP expression was further confirmed by counting the number of GFP-positive cells (Fig 3C and 4B). To assess the amount of virus produced in the presence of Selinexor, HeLa-MHVR (Fig 3D) and A549-MHVR (Fig 4C) cells were treated with different concentration of the drug and infected with MHVE-GFP. Again, like L2 cells, we observed significantly enhanced virus production in both the human cell lines when pretreated with Selinexor between 1 and 0.01 µM concentration (Fig 3D and 4C).

**Figure 3:**
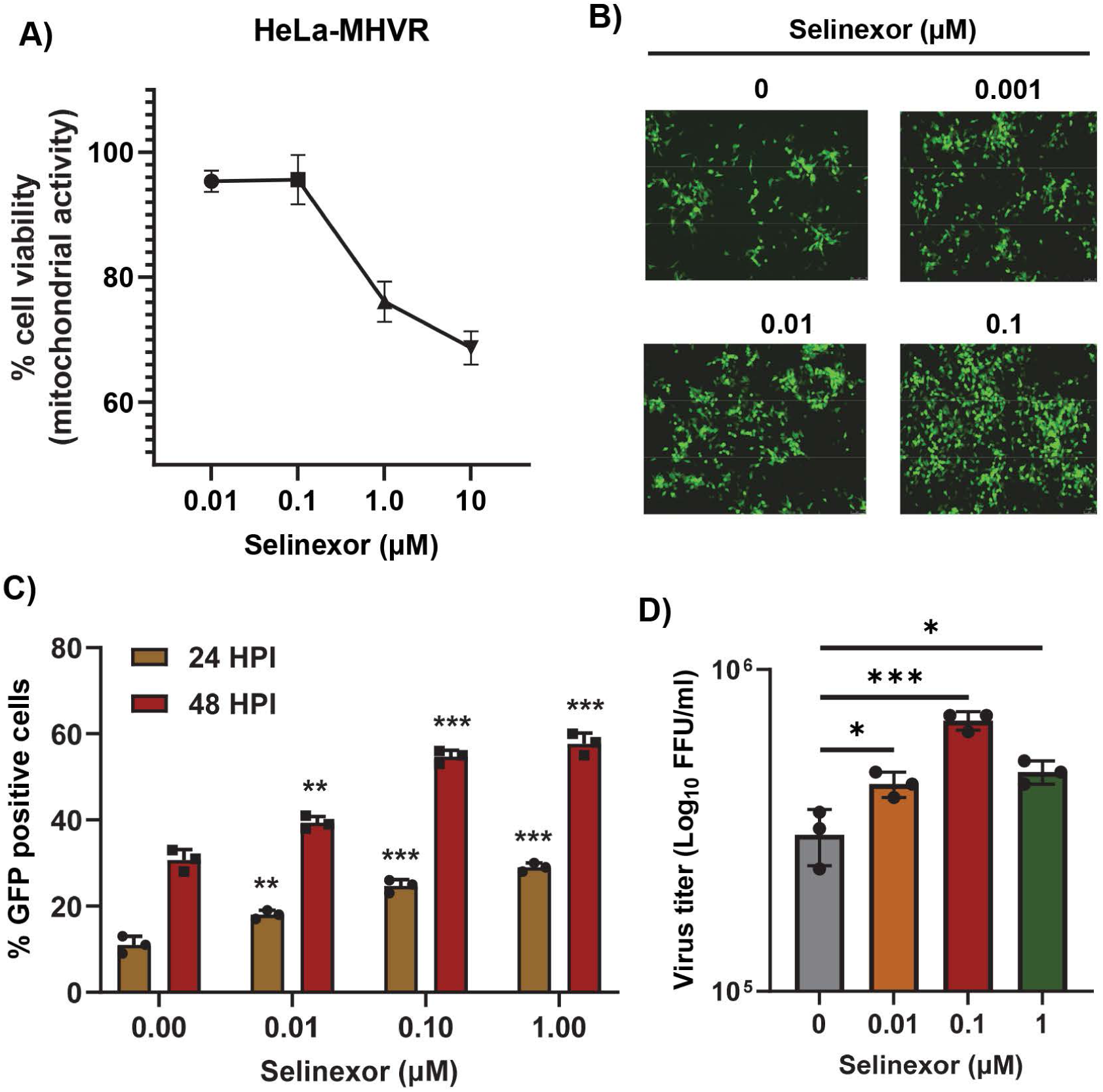
Selinexor enhances the replication of MHV in human HeLa-MHVR cells. A) Effect of Selinexor on the viability of HeLa-MHVR cells. Cells were plated in 96 well plates and treated with different concentration of Selinexor and cell viability was measured after 48h. B-D) Effect of Selinexor on MHV virus replication in HeLa-MHVR cells. Cells were plated in 24 well plates, treated with indicated concentration of Selinexor for 1h and infected with rA59/S_MHV-2_-EGFP or MHVE-GFP virus with a MOI of 0.01. B) Fluorescence images were taken 24h post infection with rA59/S_MHV-2_-EGFP; C) percent of GFP positive cells at 24h and 48h post infection with rA59/S_MHV-2_-EGFP were counted using Countess II cell counter; D) Total number of viruses in the cells and supernatant was determined by plaque assay 24h post infection of cells with MHVE-GFP. Data represents ± SD and n=3. Statistically significant differences in comparison to control are indicated. ^ns^ *P* > 0.05, * *P* < 0.05, ** *P* < 0.01, *** *P* < 0.001, **** *P* < 0.0001.

**Figure 4:**
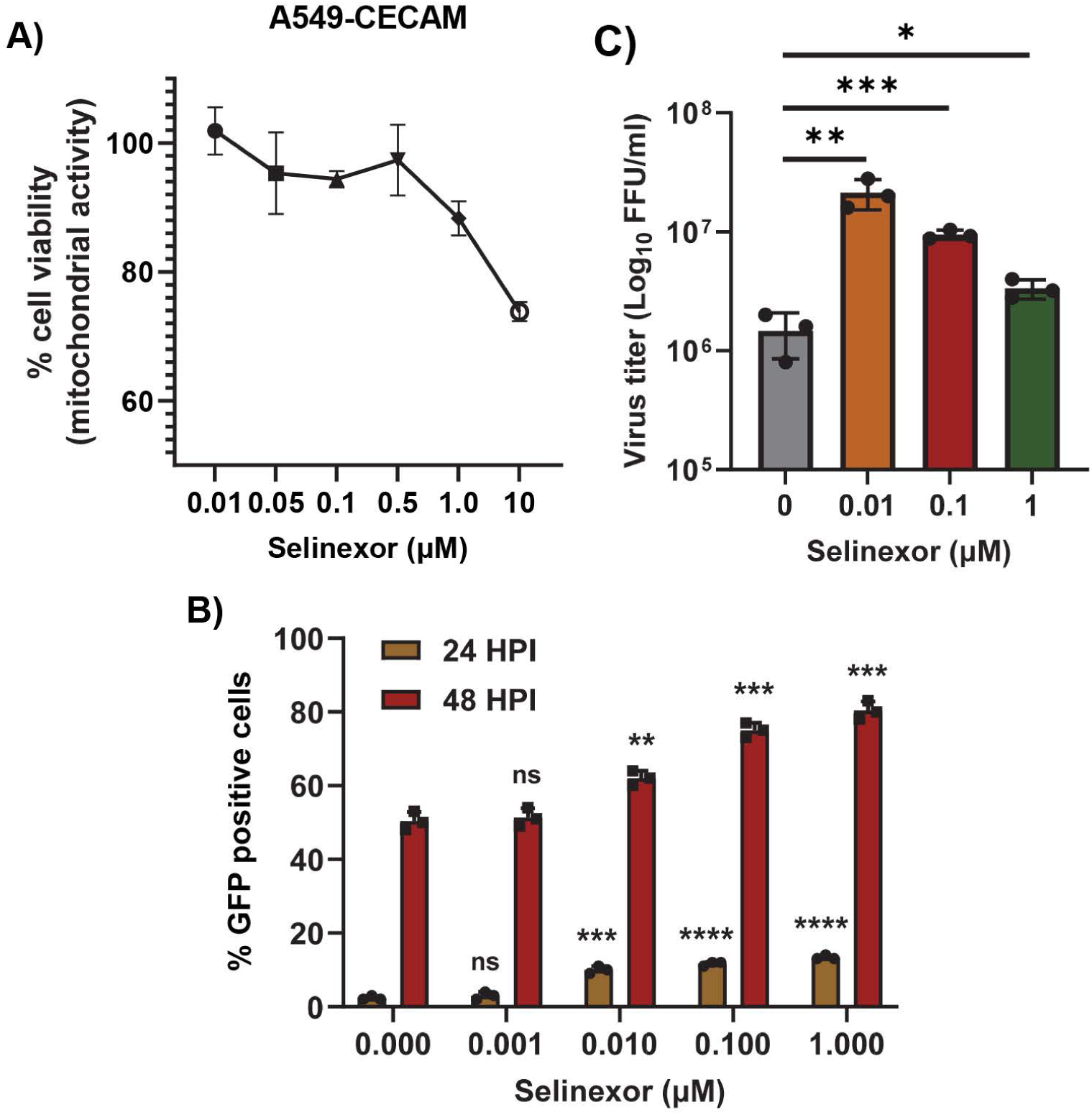
Selinexor enhances the replication of MHV in human A549-MHVR cells. A) Effect of Selinexor on the viability of A549-MHVR cells. Cells were plated in 96 well plates and treated with different concentration of Selinexor and cell viability was measured after 48h. B-C) Effect of Selinexor on MHV virus replication in A549-MHVR cells. Cells were plated in 24 well plates, treated with indicated concentration of Selinexor for 1h and infected with rA59/S_MHV-2_-EGFP or MHVE-GFP virus with a MOI of 0.01. B) percent of GFP positive cells at 24h and 48h post infection with rA59/S_MHV-2_-EGFP were counted using Countess II cell counter; C) Total number of viruses in the cells and supernatant was determined by plaque assay 24h post infection of cells with MHVE-GFP. Data represents ± SD and n=3. Statistically significant differences in comparison to control are indicated. ^ns^ *P* > 0.05, * *P* < 0.05, ** *P* < 0.01, *** *P* < 0.001.

### XPO1 knockdown enhances coronavirus gene expression and replication

Since inhibition of the XPO1-mediated nuclear export pathway using Selinexor enhanced coronavirus replication, we further extended this observation by direct knockdown of XPO1 using siRNA. After transfection of XPO1 siRNA or a non-targeting control siRNA (NT-siRNA) in A549-MHVR cells, the cells were infected with different MOIs of the GFP expressing MHV, rA59/S_MHV-2_-EGFP or MHVE-GFP. An increase in GFP-expressing cells was observed only in the XPO1 knockdown cells (Fig 5A). Furthermore, the enhanced number of GFP-positive cells in the XPO1 knockdown cells compared to controls was confirmed by counting the number of GFP-positive cells (Fig 5C). The level of XPO1 protein knockdown using siRNA was confirmed by western blot analysis (Fig 5B). To quantify the number of progeny virions in the infected cells and supernatant, the cells were infected with MHVE-GFP and samples were collected 24h post-infection. Virus titration shows that virus production significantly increased in the XPO1 knockdown cells compared to the NT-siRNA control or cells infected with the virus alone (Fig 5D), indicating that the effect of Selinexor is mediated by XPO1.

**Figure 5:**
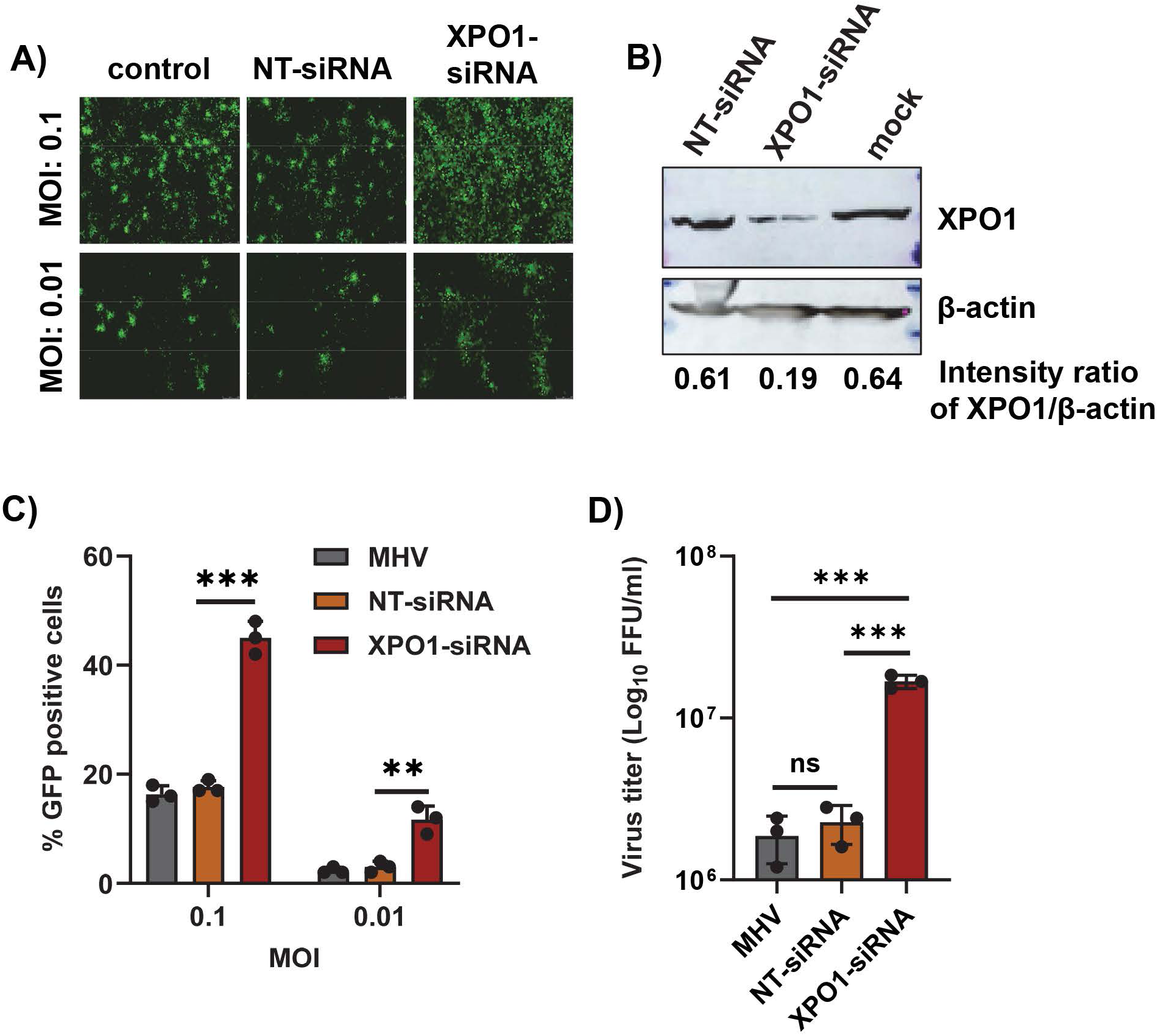
XPO1 knock down enhances the replication of MHV in human A549-CECAM cells. Cells were plated in 24 well plates, transfected with control non-targeting siRNA (NT-siRNA) or XPO1 siRNA for 48h and infected with rA59/S_MHV-2_-EGFP or MHVE-GFP for another 24h. A) Images were taken using a fluorescence microscope 24h post infection with MHVE-GFP; B) Knock down of XPO1 was confirmed with Western blot analysis using anti-XPO1 antibody. Actin was used as total protein loading control. C) percent of GFP positive cells after infection with rA59/S_MHV-2_-EGFP were counted using Countess II cell counter. D) Total number of viruses in the cells and supernatant was determined by plaque assay 24h post infection of cells with MHVE-GFP. Data represents ± SD and n=3 or 4. Statistically significant differences in comparison to control are indicated. ^ns^ *P* > 0.05, * *P* < 0.05, ** *P* < 0.01, *** *P* < 0.001.

### Effect of Selinexor on SARS-CoV-2 replication

We next tested whether Selinexor enhances the replication of SARS-CoV-2 in human cells. We used a human A549 cell line expressing the ACE2 receptor (A549^ACE2^) and a SARS-CoV-2 virus expressing GFP (SARS-CoV-2-GFP) (32). The A549^ACE2^ cell line showed a similar level of sensitivity to different concentrations of Selinexor like A549-MHVR (data not shown). To assess whether Selinexor enhances SARS-CoV-2 gene expression and replication, A549^ACE2^ cells were first pretreated with 0.01µM of Selinexor for one hour and infected with SARS-CoV-2-GFP. After 24h, cells were collected and fixed for counting the number of GFP-positive cells. A significantly increased number of GFP-positive cells were observed in the treated infected cells compared to the untreated cells (Fig 6A). We also observed a modest but significant increase in virus titer in A549^ACE2^cells (Fig 6B). The effect of Selinexor on SARS-CoV-2 was further tested using an infectious recombinant SARS-CoV-2 (rSARS-CoV-2) virus (33–35). Like the parental SARS-CoV-2 this recombinant virus titer was also increased in human A549 cells in the presence of Selinexor (Fig 7A). However, in Vero and human Calu3 cells no significant increase or decrease in virus titer was observed when treated with Selinexor (Fig 7B and 7C). These results indicate that, unlike MHV, the effect of Selinexor on the replication of SARS-CoV-2 is cell type dependent.

**Figure 6:**
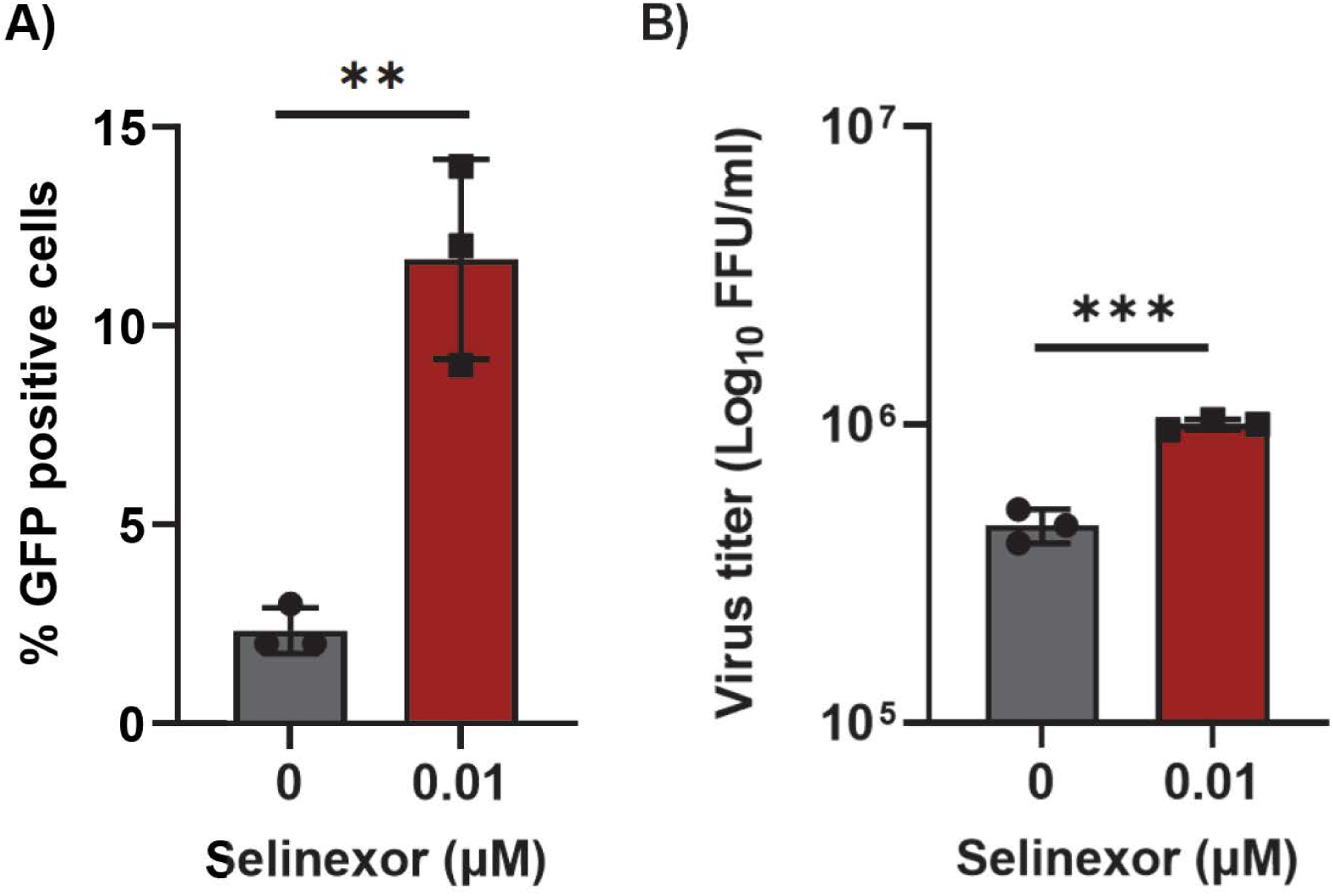
Selinexor enhances the replication of SARS-CoV-2 in human A549^ACE2^ cells. Cells were plated in 12 well plates, treated with indicated concentration of Selinexor for 1h and infected with SARS-CoV-2 virus. A) percent of GFP positive cells were counted using Countess II cell counter after fixation of cells. B) Total number of progeny virus formation was measured by plaque assay using Vero E6 cells. Data represents ± SD and n=3 or 4. Statistically significant differences in comparison to control are indicated. ^ns^ *P* > 0.05, * *P* < 0.05, ** *P* < 0.01, *** *P* < 0.001.

**Figure 7:**
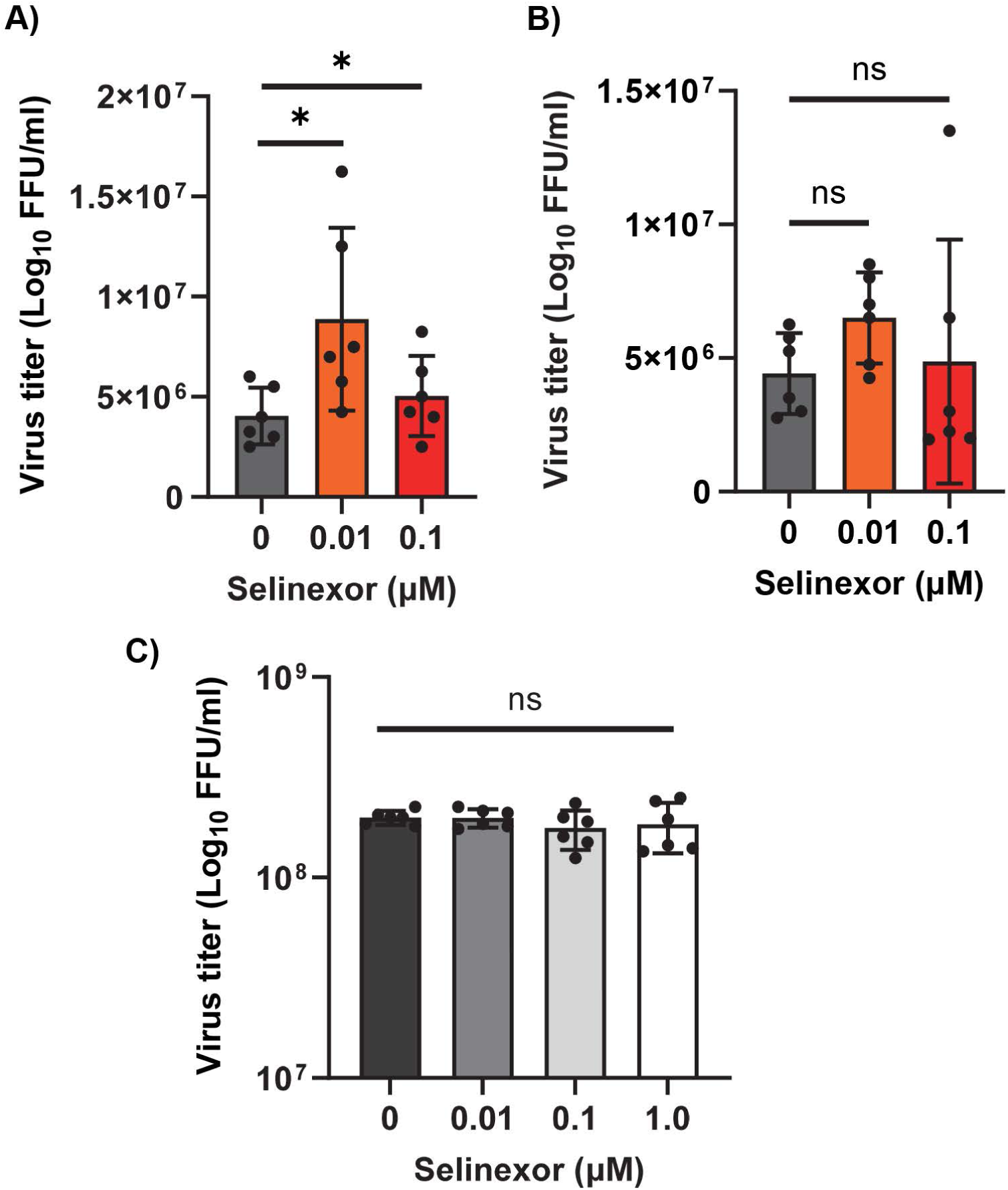
Effect of Selinexor on the replication of rSARS-CoV-2 in different cell types. A) Human A549^A2T2^ cells, B) Vero E6 cells, and C) Human Calu3 cells were plated in 12 well plates, treated with indicated concentration of Selinexor for 1h and infected with rSARS-CoV-2 virus with a MOI of 0.1. Total number of progeny virus formation was measured by plaque assay using Vero E6 cells. Data represents ± SD and n=3 or 4. Statistically significant differences in comparison to control are indicated. ^ns^ *P* > 0.05, * *P* < 0.05.

## Discussion

The nucleocytoplasmic transport process mediated by XPO1 plays a vital role in the export of hundreds of proteins from the nucleus. The proteins are involved in diverse cellular processes such as cell proliferation, cell cycle progression, and apoptosis (1, 5). Thus, the XPO1-mediated export pathway is targeted by viruses at various stages of their lifecycle to regulate cellular proteins and the appropriate localization of viral proteins (6, 36). Apart from viruses, this nuclear export pathway is also crucial for anticancer therapy due to the export of tumor suppressor proteins by XPO1 to the cytoplasm (7, 10). Therefore, XPO1 inhibitors are developed as potential antiviral and anticancer agents. The cysteine residue within the hydrophobic nuclear export sequence (NES)-binding region at position 528 is the prime target for most XPO1 inhibitors, including leptomycin B (LMB). LMB isolated from Streptomyces was the first specific inhibitor of XPO1 (37). However, the clinical development was discontinued due to severe cell toxicity (38). The irreversible binding of LMB with CRM1 caused long-term inhibition of CRM1-mediated nuclear export, and possible other off-target activity resulting in cellular toxicity (38). Synthetic derivatives of LMB with less toxicity due to the reversible binding with CRM1 have been clinically tested in human. The FDA has approved one such XPO1/CRM1 inhibitor called Selinexor for treating hematological cancers (11). XPO1 inhibitors have shown antiviral activity against many viruses, such as influenza, RSV, and recently SARS-CoV-2 (13, 25, 36, 39, 40).

We reported that, unlike RNA viruses, LMB or Selinexor enhances the replication of oncolytic MYXV, a leporipoxvirus developed for cancer treatment (26). In human cancer cells, Selinexor enhanced MYXV replication only at a low concentration that had minimal or no toxicity to the cells. However, a higher concentration of Selinexor that caused cellular toxicity also reduced virus replication. Based on our observation that nuclear export inhibitors, including Selinexor, can enhance cytoplasmic replication of a poxvirus and a study showing that Selinexor inhibits SARS-CoV-2 replication, we first tested the effect of different nuclear export inhibitors, including Selinexor, on the replication of a mouse coronavirus MHV using murine L2 cells. To our surprise, we observed that pretreatment of L2 cells with different concentrations of these inhibitors that had minimal or no toxicity to the cells significantly increased the replication of MHV. This observation was further confirmed using human HeLa and A549 cells expressing the MHV receptor murine CECAM1(30). Again, increased reporter GFP expression and MHV replication were observed when cells were pretreated with Selinexor concentrations that had minimal or no effect on the cell viability. These results confirm that the impact of Selinexor on MHV replication is independent of the cell type. Since XPO1 is the only known cellular target of Selinexor, we previously reported that XPO1 knockdown using siRNA enhanced the replication of MYXV in human cancer cells (26). Here, again, we confirmed that XPO1/CRM1 knockdown also significantly enhanced the replication of MHV in both A549-MHVR and HeLa-MHVR cell lines. Both poxviruses and coronaviruses replicate in the cytoplasm of infected cells. The observations that inhibition of XPO1-mediated nuclear export pathway can enhance the replication of both the viruses suggest the importance of this pathway for future studies to identify the host and viral proteins involved in this process. These results encouraged us to test whether the optimized lower concentration of Selinexor enhances SARS-CoV-2 replication in different cell types. However, unlike MHV, Selinexor-mediated enhanced SARS-CoV-2 replication was observed only in selected cell lines and had no significant effect on virus replication in other cell lines, such as Vero-E6 and Calu3. This observation is in contrast to a previous study by Kashyap et al. showing that in Vero E6 cells, Selinexor treatment reduced the SARS-CoV-2 plaque number and virus titer by more than two logs with >100nM Selinexor (25). In our study we did not observe any reduction in virus titer with the tested concentration of Selinexor in any of the cell lines that we tested with both MHV and SARS-CoV-2. The Selinexor concentration we used in our study had no or very minimal toxicity to the cells. However, at higher concentration of Selinexor (>100nM), we observed enhanced toxicity to the cells and can inhibit virus replication. We observed the cell type specific effect of selinexor on myxoma virus, where selinexor did not affect virus replication in rabbit RK13 cells and Vero-E6 cells (data not shown). Therefore, we can speculate that different cell types might respond to Selinexor differently against these tested viruses. Inhibition of XPO1-mediated nuclear export pathway results in several changes in the cells, such as cell cycle arrest, and modulation of cellular key signaling pathways regulating the expression of proinflammatory cytokines. Depending on the cell type, these cellular changes might impact the virus replication (41).

Selinexor is currently approved for treating selected hematological malignancies such as multiple myeloma and diffuse large B-cell lymphoma (11, 42). Multiple clinical trials are presently undergoing to use Selinexor as a monotherapy or in combination with other treatments against diverse types of malignancies (43, 44). Our results indicate that Selinexor mediated inhibition of nuclear export pathway might have different effect on virus replication. Future studies should focus on how Selinexor and nuclear transportation pathway regulate the replication of coronavirus.

## Materials and methods

### Biosafety

SARS-CoV-2-GFP virus infections and virus manipulations were conducted at biosafety level 3 (BSL-3) and rSARS-CoV-2 Δ3a/Δ7b virus experiments were conducted at BSL-2+ level in the Biodesign Institute at ASU using appropriate and IBC approved personal protective equipment and protocols.

### Cell lines

L2 murine fibroblast cell line and African green monkey kidney Vero cells E6 (obtained from ATCC) were cultured and maintained using Dulbecco’s modified Eagle’s medium (DMEM) supplemented with 10% FBS, 100 U/ml of penicillin, and 100 µg/ml streptomycin. Calu-3 cells (ATCC: HTB-55) were cultured and maintained using Eagle’s Minimum Essential Medium (EMEM) supplemented with 10% FBS, 100 U/ml of penicillin, and 100 µg/ml streptomycin. HeLa-MHVR cell line (human HeLa cell line expressing mouse coronavirus receptor mCECAM1) was provided by Tom Gallagher at Loyola University and maintained using DMEM media with 10% FBS, HEPES buffer, 100 U/ml of penicillin, 100 µg/ml streptomycin, MEM non-essential amino acids, and sodium pyruvate. Human A549 cells expressing ACE2 and TMPRSS2 (ACE2plusC3-A549/ACE2/TMPRSS2) was purchased from ATCC (CRL-3560) and maintained with DMEM media supplemented with 10% FBS, 100 U/ml of penicillin, and 100 µg/ml streptomycin. A549-MHVR (human A549 cell line expressing mouse coronavirus receptor mCECAM1) and A549^ACE2^ (human A549 cell line expressing ACE2 receptor) was provided by Susan Weiss at the Perelman School of Medicine at the University of Pennsylvania and maintained with RPMI1640 media supplemented with 10% FBS, 100 U/ml of penicillin, and 100 µg/ml streptomycin (31).

### Viruses and viral replication assay

Mouse hepatitis coronavirus MHV A59 that expresses GFP was used for all the assays. rA59/S_MHV-2_-EGFP is a recombinant MHV A59 virus that expresses the spike of MHV-2 in place of the WT protein and GFP (27–29). MHVE-GFP was constructed using a MHV A59 reverse genetics system essentially as previously described (45–48). The coding sequence for the MHV envelope (E) protein was fused to the GFP gene with an intervening tetra glycine linker. The construct was cloned into ORF 4a/b in the G subclone of the MHV infectious clone using Sbfl and EcoRV restriction sites. The construct was designed to maintain the transcription regulatory sequences for both ORF 4 and ORF5. Virus was recovered after assembling the full-length genomic cDNA and transcription of full-length genomic RNA as previously described (45–48). Following electroporation into baby hamster kidney grown in 17CI1 mouse cells virus was recovered and passage 1 stock was grown in mouse 17Cl1 cells and titered in L2 cells by plaque assay. The SARS-CoV-2-GFP virus was provided by Ralph Baric at the University of North Carolina Chapel Hill (32).

Viral titers in different cell lines were determined using a viral replication assay. The cells were seeded in 24 well plate (2×10^5^ cells/well). The next day, the cells were treated for 1h with different concentrations of nuclear export inhibitors diluted in the appropriate media used for growth of the specific cell lines. Virus was added to the cells (volume calculated based on different multiplicities of infection and incubated for 1h at 37°C in the presence of the inhibitors. After 1h, the unbound virus was removed, cells were washed with DPBS (Dulbecco’s phosphate-buffered saline) and media with inhibitors was added for further incubation. Both cells and media were collected at different time points and stored at -80°C freezer until processing. Afterwards, different dilutions were prepared in the appropriate media and plated on cell lines and fluorescent foci were counted after 24h using a fluorescent microscope. All assays and dilutions were performed in triplicate. For SARS-CoV-2, all infections and virus manipulations were conducted at biosafety level 3 (BSL-3) in the Biodesign Institute using appropriate and IBC-approved personal protective equipment and protocols. For plaque assay, samples were serially diluted 10-fold and absorbed on Vero cells at 37°C for 1h. Cells were overlaid with media plus 0.7% agarose and incubated for two days at 37°C. Cells were fixed with 4% paraformaldehyde and subsequently stained with 1% crystal violet for counting the plaques.

### Reagents and Antibodies

Rabbit polyclonal antibodies for XPO1, ACE-2 and mouse monoclonal antibody against β-actin were purchased from Thermo Fisher Scientific. HRP-conjugated goat anti-rabbit and anti-mouse IgG antibodies were purchased from Jackson Immuno Research Laboratories. Selinexor (KPT330) was purchased from Apex Bio (Tokyo, Japan). Leptomycin B, Ratjadone A, and Anguinomycin A purchased from Santa Cruz Biotechnology.

### siRNA transfection

ON-TARGETplus SMART pool siRNAs for exportin 1/XPO1 and a non-targeting control (NT siRNA) were purchased from Dharmacon (Horizon Discovery). In 24 well plate cells were seeded with 40–50% confluence, left overnight for adherence, and then transfected with siRNAs (40 nM) using Lipofectamine RNAiMAX (Invitrogen) transfection reagent. After 48 h of transfection, the cells were infected with different MOIs of virus for 1h, washed to remove the unbound virus, and incubated with complete media. At the indicated time points, cells were either observed by fluorescence microscopy to monitor and record the expression of fluorescent proteins or harvested and processed for titration of progeny virions. For detection of proteins, Western blot analysis was performed from the total proteins as described before (49).

### Cell viability assay

To assess the viability of different cell lines after Selinexor treatment, 10,000 cells were seeded into each well of a 96-well plate. The next day, cells were treated with different concentrations of Selinexor. A minimum of four to five wells were used for each treatment condition, and untreated cells (mock) served as controls. Cell viability at different time points was assessed using MTS reagents (Promega) according to manufacturer instructions.

### Statistical analysis

Statistical analyses were performed using GraphPad Prism software. Values are represented as mean ± SD for at least two or three independent experiments. ANOVA and t-test (when only two groups were compared) were used to determine the significance. *P* values are reported as follows: no significant (ns) *P* > 0.05, * *P* < 0.05, ** *P* < 0.01, *** *P* < 0.001, **** *P* < 0.0001.

## Acknowledgements

We thank Tom Gallagher (Loyola University) for the HeLa-MHVR cells; Susan Weiss (Perelman School of Medicine at the University of Pennsylvania) for the A549-MHVR cells, A549^ACE2^ cells and rA59/S_MHV-2_-EGFP virus; Ralph Baric (University of North Carolina at Chapel Hill) for the SARS-CoV-2-GFP virus; Luis Martinez-Sobrido at the Texas Biomedical Research Institute, San Antonio, TX for rSARS-CoV-2 Δ3a/Δ7b virus; and Efrem Lim and Emily Kaelin (Biodesign Institute, Arizona State University) for performing SARS-CoV-2 qRT-PCR assay.

## Funding Statement

This research was supported by grants from the National Institute of Health (NIH) USA R01 AI080607 and R21 AI163910 to M.M.R.; an Arizona Biomedical Research Center (ABRC) Investigator Award RFGA2022-010-22 to M.M.R.; an Arizona State University (Tempe, Arizona, USA) start-up grant to M.M.R.; a Mercatus Center Fast Grant (G09202-300) to G.M., M.M.R, and B.G.H.; NIH R01DK048370-26 Subaward VUMC95054 to B.G.H.; and Tohono O’Odham Nation Award GR40818 to B.G.H.

The funders had no role in the study design, data collection and interpretation, or decision to submit the work for publication.

## Notes

### Competing Interest Statement

The authors have declared no competing interest.

### Summary of Updates

This version of the manuscript has been revised to update the title, abstract, results, discussion and figures to clarify the results.

